# Learning about neurodiversity from parents – auditory gestalt perception of prelinguistic vocalisations

**DOI:** 10.1101/2023.03.13.532450

**Authors:** Dajie Zhang, Sigrun Lang, Bernd Wilken, Christa Einspieler, Jeffrey L. Neul, Sven Bölte, Daniel Holzinger, Michael Freilinger, Luise Poustka, Jeff Sigafoos, Peter B. Marschik

## Abstract

**Background:** Infants with Rett syndrome (RTT) may have subtle anomalies in their prelinguistic vocalisations but the detection of these is difficult, since their conspicuous vocalisations are often interspersed with inconspicuous ones.

**Aims and methods:** Extending a previous study with predominantly non-parents, the present study sampled parents of children with RTT and aimed to examine their gestalt perception of prelinguistic vocalisations.

**Methods and procedure:** Parents (n = 76) of female children with RTT listened to vocalisation recordings from RTT and typically developing (TD) infants, including an inconspicuous vocalisation from a RTT girl. For each recording, parents indicated if the vocalisation was produced by a RTT or a TD child.

**Results:** Overall correct to incorrect identification rate was 2:1, which was comparable to that of the previous study. Intriguingly, parents of RTT children seemed to be sensitive to features characterising the vocalisations of RTT infants, which has especially influenced their perception of the inconspicuous vocalisation from a RTT girl.

**Conclusions and implications:** These results invite further research on the potential characterising differences between vocalisations from TD infants and infants with divergent neurodevelopment.

**What this paper adds?:** Previous studies suggested that parents’ observations of their children’s behaviour are insightful and could aid clinical diagnosis. There is evidence that non-parents also seem to be sensitive to typical versus atypical characteristics in infant development. As normal and divergent developmental behaviours are often overlapping with each other, detecting deviant development is often difficult. For example, atypical vocalisations of infants later diagnosed with Rett syndrome (RTT) are often interspersed with their more typical and inconspicuous vocalisations. Can we learn extras from parents about divergences in prelinguistic vocalisations? The current study extended previous research and focused on the auditory gestalt perception of parents. We found parents of children with RTT were sensitive to the characterising differences between vocalisations from RTT and typically developing (TD) infants. They differentiated RTT vocalisations from TD vocalisations, even the RTT vocalisation was benchmarked as inconspicuous by speech-language experts. The characterising features that point to RTT, which seem to be perceptible to parents, might be more than the conspicuousness that could be readily classified by experts. What we have learned from parents’ perceptions motivates further research on the potential characterising features in prelinguistic vocalisations from different infants, especially in vocalisations that sound inconspicuous to experts and professionals, which may help to refine our understandings of diverse vocalisation patterns on the one hand, and to identify infants with neurodevelopmental divergences on the other hand.

## 1. Introduction

Rett syndrome (RTT) is a rare neurogenetic condition predominantly caused by mutations on the X-linked *MECP2* gene (Amir et al., 1999; Neul et al., 2008). Over time, our knowledge of RTT has advanced not only on its aetiology (Collins & Neul, 2022), clinical phenotypes (Ivy & Standridge, 2021) and treatments (Fu et al., 2020; Sigafoos et al., 2019), but also increasingly on its prodromal and pre-diagnostic development (Cosentino et al., 2019; Einspieler & Marschik, 2019). A period of apparently typical initial development was once a diagnostic criterion for RTT, this has been replaced by the growing acknowledgment that in development of infants later diagnosed with RTT, early subtle signs and anomalies present before the more overt symptoms of developmental regression emerge (Marschik et al., 2014; Marschik et al., 2013; Neul et al., 2010).

Indeed, retrospective and prospective studies indicate that from birth onwards, infants later diagnosed with RTT frequently follow altered developmental pathways and experience delays in achieving developmental milestones (Marschik et al., 2018; Neul et al., 2014). Disturbances in breathing, sleeping, and mood regulation patterns, for example, are often observed during early development (Buchanan et al., 2019; Tarquinio et al., 2018; Veatch et al., 2021). Abnormal neuromotor functions, such as differences in general movement patterns (Einspieler et al., 2014) and asymmetric eye-blinking (Einspieler et al., 2005) have been documented during their very first months of life in infants later diagnosed with RTT. Qualitative anomalies have also been reported in early speech-language development, such as production of vocalisations on ingressive airstream or with breathy voice characteristics (Marschik et al., 2010; Marschik, Pini, et al., 2012). These atypical vocalisations are detectable by human listeners (Marschik, Einspieler, et al., 2012) as well as by acoustic analyses using computer-based approaches (Pokorny et al., 2018; Pokorny et al., 2022). However, because the atypical vocalisations are often interspersed with more typical vocalisations in infants with RTT(Marschik et al., 2009; Marschik, Pini, et al., 2012), it can make their accurate detection challenging.

In a previous study by our research group, we assessed different participants’ auditory gestalt perception of typical versus atypical prelinguistic vocalisations (Marschik, Einspieler, et al., 2012). In that study, seven vocalisation sequences — either from typically developing (TD) infants or infants with RTT — were used as stimuli. A group of speech-language experts first benchmarked these vocalisation sequences: a sequence was defined by experts as inconspicuous if it represented age specific normal vocalisations, or conspicuous if it contained deviations from typical verbal behaviour (e.g., inspiratory character in the vocalisation; Marschik, Pini, et al., 2012). Among these seven stimuli, three vocalisations were from RTT girls and benchmarked as conspicuous, and another three were from TD girls and rated as inconspicuous. Additionally, a vocalisation from a RTT girl, which was benchmarked by experts as inconspicuous, was also included. The task of the 400 participants was to classify each vocalisation as either conspicuous or inconspicuous. The results showed that in most cases the judgements of the participants were consistent with the experts’ benchmarking, including correctly classifying the inconspicuous vocalisation produced by the RTT infant as normal. Regarding the differences between participants, the professional listeners (i.e., a subgroup of speech-language therapists, clinical linguists, and phoniatricians; n = 61) had higher accuracy rates on the tasks than the naïve listeners (i.e., college students in linguistics and medicine; n = 339). It might be worthy to mention that the vast majority of the naïve listeners were in their 20s and had no child of their own. The participants who were older than 30 years (39 out of 400) and who were parents (23 out of 400) mostly belonged to the group of speech-language professionals.

Parents are often the first to notice peculiarities in their children’s development. They have been reported to be sensitive to a multitude of early signs in the sensory-motor, speech-language, social-communicative, and neuropsychiatric domains (Bailey et al., 2003; Glascoe, 1997; Havdahl et al., 2017; Marschik, 2014; Solgi et al., 2022; Zhang et al., 2017; Zuckerman et al., 2015). With intensive and direct daily interactions with their children, parents may notice early signs of neurodevelopmental conditions that could escape the notice of professionals during episodic clinical assessments (Wallisch et al., 2020). Parental input may thus provide a unique and additional perspective which may help to complement and refine our understandings of attributes and phenomena that point to a developmental condition, which may, in turn, improve the identification and treatment of the condition.

Along these lines, the present study sought to extend previous research (Marschik, Einspieler, et al., 2012) by recruiting parents of children with RTT and exploring their auditory gestalt perception of different pre-linguistic vocalisations. In particular, we asked (a) whether parents of RTT children were able to identify the origins of vocalisations produced by either RTT or TD children; (b) whether and how the conspicuous or inconspicuous characters of the vocalisations influenced the parents’ perceptions; (c) whether the age period within which the vocalisations were produced had an effect on the parents’ performance; and (d) whether performances varied across different parents.

## 2. Materials and method

### 2.1. Participants

Participants were 76 parents (47 mothers and 29 fathers) of 59 families with a female child diagnosed with RTT. For 17 families, both parents of the child participated. In the remaining 42 families, only the mother (n = 30) or the father (n = 12) participated. All participants were members of the German Rett Parents Support Group (Rett Deutschland e.V.). Participants were recruited at a national family meeting (n = 35) and three regional meetings in 2022. The three regional meetings were in North Rhine-Westphalia (n =26), Lower Saxony (n = 8), and Saarland/Rhineland (n = 7). Detailed information of the participants and their children with RTT are presented in Table 1. In the current sample, most children (76.7%) were diagnosed before 4 years of age, with 31 (52.5%) by 24 months. Only 15 (20.3%) were diagnosed at 60 months of age or older. Most of the children (46 out of 59) had at least one sibling.

**Table 1.** Demographical data of the participants and their children with RTT.

|  | N | Variables | Mean (SD) | Median | Min | Max | P25 | P75 |
| --- | --- | --- | --- | --- | --- | --- | --- | --- |
| All participants | 76 |  | 49.0 (11.0) | 49 | 29 | 89 | 40 | 56 |
| Mothers | 47 | Age (year) | 47.8 (10.0) | 47 | 29 | 78 | 39 | 55 |
| Fathers | 29 |  | 51.0 (12.2) | 50 | 34 | 89 | 42 | 57 |
| Children with RTT | 59 | Current Age (year) | 15.5 (12.3) | 11 | 2 | 57 | 6 | 25 |
|  |  | Diagnostic Age (month) | 43.0 (41.9) | 24 | 12 | 264 | 24 | 48 |
|  |  | Number of siblings | - | 1 | 0 | 3 | 1 | 2 |

### 2.2. Materials

A total of seven audio sequences of infant prelinguistic vocalisations were used for the current experiment. These stimuli were identical to those used in a previous study (Marschik, Einspieler, et al., 2012). The audio sequences were selected from a large audio-video data pool compiled over the past decades at the research unit iDN – interdisciplinary Developmental Neuroscience, at the Medical University of Graz, Austria. A vocalisation sequence was only chosen for the present study if it was unanimously rated by a group of seven experts in linguistic and speech-language development as either inconspicuous or conspicuous (Marschik, Einspieler, et al., 2012). Of the seven sequences A to G, three were vocalisations from typically developing girls (TD; sequences B, D, and E) and were benchmarked by the experts as inconspicuous. Another three sequences were from girls with RTT and these were rated by the experts as conspicuous (sequences A, C, and F). These latter three vocalisation sequences presented with one or more of the following characteristics: (a) inspiratory components, (b) breathy/pressed voice, and/or (c) high-pitched vocalisation. The last vocalisation (sequence G) — also from a girl with RTT — was unanimously rated as inconspicuous by the experts. The sequences A to D were recorded during the first 6 months of the infants’ life, whereas sequences E to G were recorded when the infant was from 7 to 12 months of age. The duration of the sequences ranged from 23 to 27 s with a Median of 25 s. None of the recorded vocalisations were from the parent participants’ own children.

### 2.3. Procedure

The same procedure was carried out at the four recruiting sites. At each meeting, the experimenter described the study’s background and explained that participation was anonymous and voluntary. The volunteering parents of children with RTT stayed in a quiet room to complete the experiment. Each participant received a single-page worksheet. First, the participants were invited to provide the following information on the worksheet: the kinship of the participant to the child (mother/father); the age of the participant; the diagnostic age of the child with RTT; the current age of the child; and the number of siblings for the child with RTT.

Next, the participants were informed that they were going to hear seven audio clips of infant vocalisations. They were told that some of the sequences were vocalised by girls with RTT and some by typically developing girls. They were also informed that the first four sequences were vocalised by infants between 0 and 6 months of age, and that the last three sequences were from infants aged between 7 and 12 months. Their task was explained as requiring them to listen to the audio sequences carefully and then indicate on the worksheet whether they thought that vocalisation was produced by an infant with RTT. Three response options were available for each audio clip: (a) Rather yes; (b) Rather not; or (c) I do not know. The audio sequences were played via a pair of Logitech S150 USB stereo speakers connected to the experimenter’s laptop. The volume was set at 60 dB at 2 m radius from the speakers. The participants sat around the speakers at a similar distance. No training audio example was provided.

When the participants indicated that they were ready, the experimenter played the first audio sequence. Each audio sequence was played twice. Immediately after the second playing of a sequence, the participants were asked to mark their response on the worksheet, that is indicate whether the sequence was or was not vocalised by a girl with RTT. Participants were asked to make their own independent estimation without consulting with each other. After all the participants had marked their answers for the current sequence, the next audio sequence was played. The procedure continued for each audio sequence A to G, which required a total duration of about 10 min.

### 2.4. Analysis

A participant’s response was counted as correct if it was consistent with the actual origin of the vocalisation (i.e., whether the vocalisation was made by a TD or RTT infant). The response was recorded as wrong if the estimation was opposite to the origin of the vocalisation sequence, and as ambivalent if the participant chose “I do not know”.

Statistical analyses were conducted using IBM SPSS Statistics 28.0 (SPSS, Inc.). The significance level was set at p < .05. Chi-square test was applied to examine whether participants’ responses were different for different stimuli (e.g., vocalisations from RTT or TD; vocalisations that was benchmarked by experts as conspicuous or inconspicuous, respectively). Pearson correlation coefficients were calculated to assess the association between two continuous variables (e.g., the age of the participant and the age of his/her child).

## 3. Results

### 3.1. General performance

Table 2 presents the results for the 76 parents for each of the seven audio sequences. Overall, the participants made about twice as many correct responses as wrong responses. In about 10% of the cases, they were ambivalent about the origin of the vocalisation (see Table 2, Grand total).

**Table 2.** Results of the 76 participants in the auditory experiment. A participant’s response was counted as correct if it consisted with the origin (i.e., from RTT or TD infants) of the vocalisation.

| Audio Sequences |  |  |  | Number (%) of responses |  |  |  |
| --- | --- | --- | --- | --- | --- | --- | --- |
|  | Sequence | Origin | Age (month) | Correct (%) | Wrong (%) | Ambivalent (%) |  |
|  | A | RTT | 0 – 6 | 34 (44.7) | 37 (48.7) | 5 (6.6) |  |
|  | B | TD |  | 55 (72.4) | 16 (21.1) | 5 (6.6) |  |
|  | C | RTT |  | 53 (69.7) | 12 (15.8) | 11 (14.5) |  |
|  | D | TD |  | 47 (61.8) | 22 (28.9) | 7 (9.2) |  |
|  | E | TD | 7 – 12 | 50 (65.8) | 17 (22.4) | 9 (11.8) |  |
|  | F | RTT |  | 44 (57.9) | 24 (31.6) | 8 (10.5) |  |
|  | G | RTT |  | 36 (47.4) | 31 (40.8) | 9 (11.8) |  |
| Subtotal I | A – D | - | 0 – 6 | 189 (62.2) | 87 (28.6) | 28 (9.2) | *n.s.* |
|  | E – G | - | 7 – 12 | 130 (57.0) | 72 (31.6) | 26 (11.4) |  |
| Subtotal II | A, C, F, G | RTT | - | 167 (54.9) | 104 (34.2) | 33 (10.9) | *sig.*<sup>a</sup> |
|  | B, D, E | TD | - | 152 (66.7) | 55 (24.1) | 21 (9.2) |  |
| Subtotal III | A, G | - | - | 70 (46.1) | 68 (44.7) | 14 (9.2) | *sig.*<sup>b</sup> |
|  | B – F | - | - | 249 (65.5) | 91 (23.9) | 40 (10.5) |  |
| Subtotal IV | C, F | RTT | - | 97 (63.8) | 36 (23.7) | 19 (12.5) | *n.s.* |
|  | B, D, E | TD | - | 152 (66.7) | 55 (24.1) | 21 (9.2) |  |
| Grand total | A – G | - | - | 319 (60.0) | 159 (29.9) | 54 (10.2) |  |

Key: *n.s.*, non-significant; *sig.*, significant; <sup>a</sup>, $\chi^2 (2, 76) = 7.77, p = .021$ ; <sup>b</sup>, $\chi^2 (2, 76) = 22.75, p < .0001$ .

As shown in Table 2, the participants’ performances did not differ significantly depending on whether the vocalisations were produced by younger (0-6 months) or older (7-12 months) infants (Table 2, Subtotal I). However, their performances differed significantly depending on the origins of the vocalisations. Specifically, the performance (i.e., the proportion of correct responses) was poorer for the vocalisations from children with RTT (from hereon: the RTT < TD pattern; Table 2, Subtotal II). Taking a closer look, the performance for the vocalisation sequences A and G (both originating from RTT children) were significantly poorer than the ones for the remaining five sequences B to F (from hereon: the AG < B – F pattern; Table 2, Subtotal III). The performances for the sequences B to F (including sequences originating from both RTT and TD children), however, were comparable to each other; *χ*^2^ (8, 76) = 9.18, *p =* .37 (see also Table 2, Subtotal IV).

### 3.2. Performance of the current vs. the previous study (Marschik, Einspieler, et al., 2012)

Overall, the current sample of parents performed similarly to the sample in the Marschik, Einspieler, et al. (2012) study, with a notable exception related to the sequence G, as will be reported further below.

In both studies, participants knew that some of the audio sequences were vocalised by TD infants and some by infants with RTT. In the current study, participants were asked to report whether they thought each vocalisation was from a girl with RTT; whereas in the previous study, the participants were asked to classify whether each vocalisation was inconspicuous or conspicuous. Mind the twist that the audio sequence G was vocalised by a girl with RTT, but benchmarked as inconspicuous by the experts. Accordingly, we compared the performances for the sequence G separately.

For the six sequences A to F, the performance of the current sample was 62.1% correct, 28.1% wrong, and 9.9% ambivalent. In the previous study with 400 participants, the results were nearly identical for the sequences A to F, with 62.2% correct, 26.0% wrong, and 11.8% ambivalent.

In the previous study, a subgroup of speech-language professionals (n = 61) was more accurate than the college students (n = 369) in classifying the vocalisations as either conspicuous or inconspicuous. For the six sequences A to F, the professionals made 72.7% correct responses, whereas the students had 59.2% correct.

For the sequence G, an inconspicuous vocalisation from a RTT girl, the tasks and consequently the response patterns of the participants were different between the two studies. In the current study, 47.4% of the parents indicated that the sequence G was vocalised by a RTT infant, while 40.8% responded that it was not from an infant with RTT, and 11.8% were ambivalent. In other words, a substantial proportion of the parents correctly identify that the sequence G was from an infant with RTT, even though experts rated this sequence as inconspicuous. In the previous study, where the task was to detect conspicuousness of the vocalisation, across all the 400 participants, 61.5% chose inconspicuous, 17.5% conspicuous, and 21.0% were ambivalent. Again, the professionals were more accurate (74.0%) than the students (59.8%) in identifying the sequence as inconspicuous.

### 3.3. Performances in the subgroups

The participants were different in their age, their kinship to the child (mother or father), and their child-rearing experience (i.e., had only one child vs. two or more children). The chronical age and the diagnostic age of their children were different (Table 1). We examined whether these variables were associated with the participants’ responses.

The overall performance (i.e., across all the seven audio sequences A to G) of the participants did not significantly differ between the participants’ age groups (younger vs. older; grouped by median, see Table 1), nor between mothers and fathers (see detailed data of each subgroup for each audio sequence in Table S1 in the Supplementary Materials). The participant’s age was highly correlated with the age of the child with RTT (mother with child: r(45) = .89; father with child: r(27) = .90). As such, and similarly, the participants’ overall performance also did not significantly differ from each other regarding whether they had a younger or an older child with RTT. Additionally, whether the mothers and the fathers had a child diagnosed earlier or later (grouped by median, see Table 1), or whether they had only one child or more children, also did not significantly influence the participants’ overall performance.

Note that whether the parents were younger or older did not relate to whether they had raised one child or more children. Among the younger parents (n = 36), 7 had only one child (i.e., the child with RTT), and the other 29 had two or more children (i.e., the child with RTT and at least one more child). Among the older parents (n = 40), 10 had only one child, and the remaining 30 had two or more children. In other words, the child-rearing experience did not significantly differ between the younger and the older parents. Accordingly, in the following analyses, if a difference would be found between the younger and the older parents, this would not be further interpreted regarding whether the difference was related to the parents’ child-rearing experience.

In addition, the parent’s age was correlated with the diagnostic age of the child: the younger the parent, the earlier their child was diagnosed; *χ*^2^ (2, 76) = 16.23, *p* < .001. As such, we did not report the potential effect of the child’s diagnostic age on the parent’s performance (data can be requested from the authors). Instead, we report in the following sections the effect of the participants’ own age on their performance.

#### 3.3.1. Different participants and the RTT < TD pattern

Although an inferior performance for the vocalisations from RTT than TD was found for the entire sample (Table 2, Subtotal II), the RTT < TD pattern was only significant in the fathers but not in the mothers (Table S2, Subtotal II; Supplementary Materials). The RTT < TD pattern was also comparable in the younger and the older participants (Table S2, Subtotal I; Supplementary Materials).

#### 3.3.2. Different participants and the AG < B – F pattern

Although there was an AG < B – F pattern in the entire sample (Table 2, Subtotal III), this was not equally pronounced in the participants. In particular, the AG < B – F pattern was only significant in the older mothers, *χ*^2^ (2, 24) = 19.40, *p* < .001, but not the younger mothers, nor the fathers (Table S3; Supplementary Materials).

## 4. Discussion

In this study, 76 parents of children with RTT listened to vocalisations either from TD or RTT girls. When asked to estimate the origin of the vocalisations (i.e., from RTT or not), the parents made about twice as many correct responses as incorrect identifications. This accuracy rate was comparable to the 400 participants in a previous study (Marschik, Einspieler, et al., 2012). The current participants’ performance did not differ whether the vocalisations were recorded at a younger (0 to 6 months) or older age (7 to 12 months). The participants’ overall performances did not differ between younger and older parents, nor between mothers and fathers. The participants’ child-rearing experience also did not influence their overall performance. Some significant differences for certain audio sequences were found and are discussed further below.

The participants’ task in the current study was adapted to their background, i.e., raising or having raised a daughter with RTT, and was slightly different from that used in the Marschik, Einspieler, et al. (2012) study. In the current study, the participants were asked to indicate whether they thought each vocalisation sequence was produced by an infant with RTT. In the previous study, in contrast, the participants were required to distinguish between conspicuous versus inconspicuous vocalisations. For the six “congruent” audio sequences A to F, where the conspicuous vocalisations were produced by infants with RTT, and the inconspicuous ones by TD infants, the tasks of the two studies were comparable. Indeed, the participants’ performances across the two studies were nearly identical for the congruent sequences A to F. In both studies and in most cases, the participants correctly classified RTT versus TD and conspicuous versus inconspicuous vocalisations. Their classification accuracy was higher than chance level.

The vocalisations of RTT girls vary from child to child, although some peculiar characteristics, such as inspiratory, high-pitched, or breathy voice components have frequently been reported in vocalisations from RTT girls (Marschik et al., 2010; Marschik, Pini, et al., 2012). However, infants with RTT do not constantly produce conspicuous vocalisations. Quite the opposite, their atypical or conspicuous vocalisations are often interspersed with more typical or inconspicuous vocalisations (Marschik et al., 2009; Marschik et al., 2013; Marschik, Pini, et al., 2012). Given the individual differences, and the interspersing of typical and atypical vocalisation within individuals, a clear-cut “RTT-like” or “TD-like” vocalisation pattern may not exist, and conspicuous vocalisations from RTT children may not always be apparent, especially not to the listeners with no professional training in speech-language pathology (Marschik et al., 2016).

Indeed, in the Marschik, Einspieler, et al (2012) study, a subgroup of speech-language professionals had overall higher correct identification rates than both the college students in that study and the parents in the present study. The professionals’ judgements were more consistent with the experts’ benchmarking. This suggests that the professional training and practice in speech-language development and pathology, but not necessarily the experience of having a child with a neurodevelopmental condition (as had the participants in the current study), may heighten the participants’ acuity with respect to detecting conspicuous prelinguistic vocalisations. This finding coincides with the results of another study investigating whether day-care workers could identify deviant development in young children (Zhang et al., 2019). The authors found that the day-care workers with advanced pedagogical training and practice in children with special needs were better at detecting atypical features in early development. In contrast, carers with years of experience of nursing young children, but no special pedagogical training, did not perform as well, although their performance was still above chance levels. In short, evidence suggests that specialised training and targeted practice may sharpen the individual skills in a task requiring detection of peculiarities in developmental behaviours (see also Lee et al., 2005; Reilly et al., 2015).

The result for the audio sequence G was intriguing. Importantly, the sequence G was rated as inconspicuous or normal by the experts, although it came from a RTT girl. Remarkably, about 47% of the parents identified, still, that the inconspicuous sequence G was from a RTT girl. As a contrast, for the other three inconspicuous vocalisations (all from TD infants), only about 24% of the parents assumed they were likely made by RTT infants, while 67% of the parents recognised that the vocalisations were made by TD infants (Table 2, Subtotal II). The significant estimation pattern shift suggests that, the sequence G, although being inconspicuous, might contain certain “characterising features” that make it sound “RTT-like”, which seemed to be detectable by parents of RTT children. Still, 41% of the parents thought sequence G might be produced by a TD girl, implying that the inconspicuous character of G might also be perceptible to the parents. Compared to the previous Marschik, Einspieler, et al. (2012) study, the participants were not asked to identify “who” made the vocalisation, but whether the vocalisation sounded inconspicuous or not. In that study, most participants (62%) rated G as inconspicuous, similar as they did for the other three inconspicuous sequences. We do not know, however, whether those participants would have judged differently if they were asked to identify “who” made the vocalisation. Unfortunately, only one inconspicuous vocalisation from RTT infants was included in this study. It is not clear what features the parents may have been perceptive to, which would be an interesting question for future research, especially with a higher number of inconspicuous vocalisations from infants with RTT.

In recent years, with automated signal level analysis in infant vocalisations, researchers demonstrated that vocalisations from infants with a developmental disorder or infants with typical development can be reliably distinguished (Pokorny et al., 2022). Further studies are especially needed to examine the potential subtle differences between the inconspicuous vocalisations either from children with typical development or from children with a developmental disorder, such as RTT. Vocalisations from children with neurodevelopmental divergences, even if they sound inconspicuous for the ears of speech-language experts and professionals, may bear inherent characterising features revealing their origins that are perceptible to others, such as parents. Just like once we assumed that the pre-regression development of individuals with RTT was just normal, we might, learning through the perspectives of parents, modify our understandings of inconspicuousness versus conspicuousness, not only in the speech-language domain.

It is important to bear in mind that typically developing children also produce vocalisations that sound conspicuous from time to time. For example, transient high-pitched crying in pain, pressed voice, or articulations with inspiratory airstream during pleasure bursts are observable (Marschik et al., 2022; Nathani & Oller, 2001; Oller, 2000). Could humans and computer algorithms also in those cases detect the “origin-revealing” features and correctly identify the producers of the conspicuously-sounding vocalisations? A related question could be to ask whether TD infants and infants with divergent development differ primarily in the *quantity* of conspicuous utterances they produce, or, are there reliably detectable *qualitative* differences? In any case, the origin-revealing or characterising features, if they could be defined, will critically enhance the sensitivity and specificity in early detection of divergent prelinguistic development.

One limitation of the present study was that the presenting order of the audio sequences was fixed from A to G at different recruiting sites and no audio example was provided before the formal experiment. The participants’ lower than average performance for the sequence A (45% correct vs. 49% wrong estimations, Table 2) might stem from then being unprepared and unpractised in performing the task. This result was similar in the Marschik, Einspieler, et al. (2012) study, where the stimuli presentation also directly begun with the sequence A, and its detection rate was 43% correct vs. 47% wrong. Had we randomised the order of stimuli presentation across sites, and offered warming-up examples for the participants, we might have obtained different results.

With the above discussion, the causes for the AG < B – F pattern (Table 2) may have been explained. Although the correct response rates for the sequences A and G were both lower than average, their causes might be dissimilar. The statistically significant RTT < TD pattern (Table 2) was also primarily caused by the lower estimation accuracy of A and G, but not by the origins of the vocalisations (i.e., RTT or TD) per se. Afterall, the participants’ response rates were all comparable for the remaining five sequences B to F (Table 2, Subtotal IV), including vocalisations from both RTT (C and F) and TD (B, D, and E) infants.

The overall performances between the parents (i.e., younger and older mothers, and younger and older fathers) did not significantly differ from each other. Given that the sample sizes of the subgroups were small and the number of stimuli (the audio sequences) was limited, any difference between the subgroups shall be interpreted with caution. Especially for the fathers, divided by their median age, only 13 belonged to the younger group and 16 the older group. We shall thus pool the data of the male participants in one group for discussion. It was found that the older mothers had even lower estimation accuracy for the sequences A and G, but on the other hand, higher accuracy for the other sequences B to F, making their overall performance just as good as the younger mothers and the fathers. In other words, it was the contrast between the performances for A, G and the rest sequences that was greater in the older mothers than in the other subgroups (Table S1 and S3, Supplementary Materials). Similarly, the contrast between the performances for the vocalisations from RTT and TD infants was greater in the fathers than in the mothers (Table S3; Supplementary Materials), which again, shall not be overinterpreted given that their overall performances were comparable. To draw conclusions on the performances of the subgroups, not only bigger sample sizes are required, but also a higher number of stimuli would be inevitably needed.

Our study had several shortcomings. Besides the aforementioned limited number of stimuli and the fixed order of their presentation, we could only carry out group-wise experiments instead of testing only one participant at a time. However, group-wise trials with limited number of stimuli might be the only realistic approach at such meetings with parents of children with RTT. Although we explicitly asked the participants to make their own estimations, we cannot exclude the possibility that the participants influenced each other. Especially the participants from the same family might have even stronger influences on each other. Although 76 participants do not form a large sample, it is a considerably sound sample size for conducting research with families with children of rare diseases. We therefore are cautious in interpreting the differences between the subgroups. Nevertheless, the results of the current study resembled a previous similar study with 400 participants, which is reassuring regarding the representativeness of the current sample.

## 5. Conclusion

Our results suggest that parents of children with RTT appear to be sensitive to the differences between vocalisations from RTT versus TD infants. Parents may prove to be valuable informants especially for revealing subtle yet characterising features in prelinguistic vocalisations from individuals with RTT. Mounting evidence highlights peculiarities in very early neurofunctional constructs in infants later diagnosed with a developmental condition. With input from experienced parents and trained professionals, our understanding on neurodevelopmental diversities will be enriched and refined, which will continuously optimise early detection of developmental disorders that are in part still not detected early enough.

## Acknowledgement

We sincerely thank all the families who consented to participate the study.

## CRediT authorship contribution statement

Conceptualisation: D.Z., C.E., P.B.M.; Methodology: D.Z., P.B.M.; Investigation: S.L., P.B.M.; Formal analysis: D.Z..; Resources: D.Z., C.E., P.B.M.; Project administration and Data pre-processing: S.L.; Writing—original draft: D.Z., J.S., P.B.M.; Writing—review and editing: all authors; Supervision: P.B.M.; Funding acquisition: D.Z., B.W., P.B.M. All authors have read and agreed to the final version of the manuscript.

## Data availability

Data will be made available on request

