## Supplementary Materials for "Learning about neurodiversity from parents – auditory gestalt perception of prelinguistic vocalisations"

Table S1. Number of participants’ responses that were correct (Cr), wrong (Wr), or ambivalent (Am) for each audio sequence. The median ages (mothers: 47 years; fathers: 50 years) were used to group the participants. Grey shades highlight the participants’ performance on the respective audio sequence that was inferior than average.

| **Audio Origin** |  | **Vocalisations in the first half year of life (0-6 months)** | | | | | | |  | **Vocalisations in the second half year of life (7-12 months)** | | | | | | |  |  |
| --- | --- | --- | --- | --- | --- | --- | --- | --- | --- | --- | --- | --- | --- | --- | --- | --- | --- | --- |
|  |  | **A** | | |  | **C** | | |  | **F** | | |  | **G** | | |  | **Sum** |
|  |  | **Cr** | **Wr** | **Am** |  | **Cr** | **Wr** | **Am** |  | **Cr** | **Wr** | **Am** |  | **Cr** | **Wr** | **Am** |  |  |
|  | **Mother** | **19** | **25** | **3** |  | **31** | **8** | **8** |  | **30** | **12** | **5** |  | **21** | **20** | **6** |  | **47** |
|  | younger | 11 | 9 | 3 |  | 14 | 4 | 5 |  | 14 | 5 | 4 |  | 12 | 9 | 2 |  | 23 |
|  | older | 8 | 16 | 0 |  | 17 | 4 | 3 |  | 16 | 7 | 1 |  | 9 | 11 | 4 |  | 24 |
| **RTT** | **Father** | **15** | **12** | **2** |  | **22** | **4** | **3** |  | **14** | **12** | **3** |  | **15** | **11** | **3** |  | **29** |
|  | younger | 8 | 3 | 2 |  | 9 | 3 | 1 |  | 5 | 7 | 1 |  | 6 | 6 | 1 |  | 13 |
|  | older | 7 | 9 | 0 |  | 13 | 1 | 2 |  | 9 | 5 | 2 |  | 9 | 5 | 2 |  | 16 |
|  | **Sum** | **34** | **37** | **5** |  | **53** | **12** | **11** |  | **44** | **24** | **8** |  | **36** | **31** | **9** |  | **76** |
|  |  | **B** | | |  | **D** | | |  | **E** | | |  |  |  |  |  | **Sum** |
|  |  | **Cr** | **Wr** | **Am** |  | **Cr** | **Wr** | **Am** |  | **Cr** | **Wr** | **Am** |  |  |  |  |  |  |
|  | **Mother** | **35** | **9** | **3** |  | **29** | **15** | **3** |  | **32** | **9** | **6** |  |  |  |  |  | **47** |
|  | younger | 16 | 6 | 1 |  | 15 | 7 | 1 |  | 16 | 5 | 2 |  |  |  |  |  | 23 |
|  | older | 19 | 3 | 2 |  | 14 | 8 | 2 |  | 16 | 4 | 4 |  |  |  |  |  | 24 |
| **TD** | **Father** | **20** | **7** | **2** |  | **18** | **7** | **4** |  | **18** | **8** | **3** |  |  |  |  |  | **29** |
|  | younger | 7 | 4 | 2 |  | 7 | 3 | 3 |  | 8 | 4 | 1 |  |  |  |  |  | 13 |
|  | older | 13 | 3 | 0 |  | 11 | 4 | 1 |  | 10 | 4 | 2 |  |  |  |  |  | 16 |
|  | **Sum** | **55** | **16** | **5** |  | **47** | **22** | **7** |  | **50** | **17** | **9** |  |  |  |  |  | **76** |

Table S2. Participants’ responses on vocalisations either from girls with RTT (RTT; audio sequences A, C, F, and G) or from typically developing girls (TD; audio sequences B, D, and E). The median ages (mothers: 47 years; fathers: 50 years) were used to group the participants. A participant’s response was counted as correct if it consisted with the origin of the respective audio sequence (i.e., from RTT or TD infants).

|  | Parents | | *N* | Vocalisation origin | Number (%) of responses | | | | | |  |  |  |
| --- | --- | --- | --- | --- | --- | --- | --- | --- | --- | --- | --- | --- | --- |
|  | Age | Kinship |  |  | Correct *(%)* | | Wrong *(%)* | | Ambivalent *(%)* | |  |  |  |
|  | Young | Mother | 23 | RTT | 52 | *(56.5)* | 28 | *(30.4)* | 12 | *(13.0)* |  | *-* | |
|  |  |  |  | TD | 44 | *(63.8)* | 20 | *(29.0)* | 5 | *(7.2)* |  |  |  |
|  |  | Father | 13 | RTT | 25 | *(48.1)* | 23 | *(44.2)* | 4 | *(7.7)* |  |  |  |
|  |  |  |  | TD | 25 | *(64.1)* | 9 | *(23.1)* | 5 | *(12.8)* |  |  |  |
|  | Old | Mother | 24 | RTT | 58 | *(60.4)* | 29 | *(30.2)* | 9 | *(9.4)* |  |  |  |
|  |  |  |  | TD | 49 | *(68.1)* | 15 | *(20.8)* | 8 | *(11.1)* |  |  |  |
|  |  | Father | 16 | RTT | 32 | *(50.0)* | 24 | *(37.5)* | 8 | *(12.5)* |  |  |  |
|  |  |  |  | TD | 34 | *(70.8)* | 11 | *(22.9)* | 3 | *(6.3)* |  |  |  |
| Subtotal I | Young |  | 36 | RTT | 77 | *(53.5)* | 51 | *(35.4)* | *16* | *(11.1)* |  | *n.s.* | *n.s.* |
|  |  |  |  | TD | 69 | *(63.9)* | 29 | *(26.9)* | *10* | *(9.3)* |  |  |  |
|  | Old |  | 40 | RTT | 90 | *(56.3)* | 53 | *(33.1)* | *17* | *(10.6)* |  | *^§^* |  |
|  |  |  |  | TD | 83 | *(69.2)* | 26 | *(21.7)* | *11* | *(9.2)* |  |  |  |
| Subtotal II |  | Mother | 47 | RTT | 110 | *(58.5)* | 57 | *(30.3)* | 21 | *(11.2)* |  | *n.s.* | *^†^* |
|  |  |  |  | TD | 93 | *(66.0)* | 35 | *(24.8)* | 13 | *(9.2)* |  |  |  |
|  |  | Father | 29 | RTT | 57 | *(49.1)* | 47 | *(40.5)* | 12 | *(10.3)* |  | *sig. ^a^* |  |
|  |  |  |  | TD | 59 | *(67.8)* | 20 | *(23.0)* | 8 | *(9.2)* |  |  |  |
| Subtotal III |  |  | 76 | *RTT* | 167 | *(54.9)* | 104 | *(34.2)* | 33 | *(10.9)* |  | *sig. ^b^* | |
|  |  |  |  | *TD* | 152 | *(66.7)* | 55 | *(24.1)* | 21 | *(9.2)* |  |  |  |
| Grand total | |  | 76 | *-* | 319 | *(60.0)* | 159 | *(29.9)* | 54 | *(10.2)* |  |  |  |

*Key: n.s., non-significant; sig., significant; ^a^*, *χ ^2^* (2, 29) = 7.73, *p* = .021; ^b^, *χ ^2^* (2, 76) = 7.77, *p* = .021; *^§^*, *χ ^2^* (2, 76) = 5.19, *p* = .075; *^†^*, *χ ^2^* (6, 76) = 11.46, *p* = .075

Table S3. Participants’ responses on the audio sequences A and G versus B to F. The median ages (mothers: 47 years; fathers: 50 years) were used to group the participants. A participant’s response was counted as correct if it consisted with the origin of the respective audio sequence (i.e., from RTT or TD infants).

| Parents | *N* | Audio sequences | Number (%) of responses | | | | | |  |  |
| --- | --- | --- | --- | --- | --- | --- | --- | --- | --- | --- |
|  |  |  | Correct *(%)* | | Wrong *(%)* | | Ambivalent *(%)* | |  |  |
| Young | 36 | A, G | 37 | *(51.4)* | 27 | *(37.5)* | 8 | *(11.1)* | *n.s.* | *χ ^2^* (6, 76) = 28.42, *p* < .001 |
|  |  | B – F | 111 | *(61.7)* | 48 | *(26.7)* | 21 | *(11.7)* |  |  |
| Old | 40 | A, G | 33 | *(41.3)* | 41 | *(51.3)* | 6 | *(7.5)* | *χ ^2^* (2, 76) = 24.32, *p* < .001 |  |
|  |  | B – F | 138 | *(69.0)* | 43 | *(21.5)* | 19 | *(9.5)* |  |  |
| Mother | 47 | A, G | 40 | *(42.6)* | 45 | *(47.9)* | 9 | *(9.6)* | *χ ^2^* (2, 76) = 21.12, *p* < .001 | *χ ^2^* (6, 76) = 24.67, *p* < .001 |
|  |  | B – F | 157 | *(66.8)* | 53 | *(22.6)* | 25 | *(10.6)* |  |  |
| Father | 29 | A, G | 30 | *(51.7)* | 23 | *(39.7)* | 5 | *(8.6)* | *n.s.* |  |
|  |  | B – F | 92 | *(63.4)* | 38 | *(26.2)* | 15 | *(10.3)* |  |  |
| Subtotal | 76 | A, G | 70 | *(46.1)* | 68 | *(44.7)* | 14 | *(9.2)* | *χ ^2^* (2, 76) = 22.75, *p* < .001 | |
|  | 76 | B – F | 249 | *(65.5)* | 91 | *(23.9)* | 40 | *(10.5)* |  |  |
| Grand total | 76 | A – G | 319 | *(60.0)* | 159 | *(29.9)* | 54 | *(10.2)* |  |  |
